# Advancing High-Resolution 7T Diffusion MRI: Evaluating Phase-Encoding Correction Strategies for Distortion Correction from Basic to Four-Way Acquisitions

**DOI:** 10.64898/2025.12.01.687254

**Authors:** Kurt G. Schilling, Alexander J.S. Beckett, Matthew Amandola, Erica B. Walker, David A. Feinberg, Silvia A. Bunge, An T. Vu

## Abstract

**Purpose:** High-resolution 7T diffusion MRI (dMRI) is limited by image artifacts that compromise anatomical accuracy. The purpose of this study was to systematically evaluate phase-encoding (PE) acquisition and correction strategies to determine which methods best mitigate geometric distortions and improve data reproducibility.

**Methods:** Five healthy adults were each scanned twice on a 7T MRI scanner (0.9 mm isotropic resolution), using a highly oversampled dMRI protocol with four PE directions (AP, PA, RL, LR). From this dataset, we created and processed eleven time-equivalent, 10-minute acquisitions, ranging from uncorrected single-PE data to comprehensive 4-way PE schemes. These strategies were quantitatively compared on their geometric alignment with T1-weighted images and on the scan-rescan reproducibility of DTI-derived metrics.

**Results:** (1) All distortion-corrected schemes significantly improved geometric accuracy over uncorrected data; (2) Strategies correcting with a full set of reversed-PE (2-way) diffusion weighted images (DWIs) outperformed the common approach of using only a single reversed *b*=0 image; and (3) a 4-way PE acquisition consistently provided the highest image fidelity and reproducibility. The optimized acquisition enabled high-quality reconstruction of both long-range and fine-scale superficial white matter pathways.

**Conclusion:** For high-resolution 7T dMRI, multi-PE acquisition is essential to achieve accurate geometry and stable microstructural estimates (i.e., less residual EPI distortion and better scan-rescan agreement). A 4-way PE scheme provides the most accurate and reproducible results for microstructural and connectivity modeling.

**Data statement:** Data will be made available in BIDS format upon acceptance of the manuscript. To be updated with DOI.

## 1. Introduction

High field 7 Tesla (7T) diffusion MRI presents a powerful opportunity to map the human brain’s structural connections and tissue microstructure in unprecedented detail. The significant gain in signal-to-noise ratio (SNR) can be traded for submillimeter spatial resolution or advanced diffusion contrast, enabling in vivo mapping of highly specific tissue microstructural measures and resolving fine-scale structural connections with tractography. Recent advances in hardware and sequence development, including high performance gradients, multi-channel transmit/receive arrays, and accelerated acquisitions, have now made whole-brain, high-resolution studies feasible with reasonable scan times. While these breakthroughs open avenues for neuroscience research, the high field strength also exacerbates image artifacts that can compromise anatomical accuracy, modeling, and reproducibility.

A primary challenge at 7T is the increased severity of susceptibility-induced distortions inherent to echo planar imaging (EPI), the standard acquisition for diffusion MRI. Local field inhomogeneities at air–tissue interfaces, particularly near the sinuses and ear canals, produce large frequency offsets that displace signal in the phase-encode (PE) direction. At high fields, the differences in magnetic susceptibility at air-tissue interfaces (e.g., in the sinuses and temporal lobes) are magnified, leading to large off-resonance frequency shifts that cause geometric distortion along the phase-encode direction and intensity artifacts - signal dropout (through-plane dephasing) and pile-up (voxel compression).

These distortions are critical to address, because they significantly warp anatomical structures, misalign diffusion data with anatomical scans, broaden point-spread function (in areas with signal pile-up), and ultimately impair the accuracy of tractography and other diffusion-derived maps. Accurate image geometry is fundamental to all neuroimaging analysis and interpretation. The ability to localize findings - whether in white or gray matter - is necessary for aligning data with other imaging modalities, comparing results across subjects, and ultimately answering the question of “where” anatomical or functional changes occur. Although this principle is true for all analyses, it is especially critical for fiber tractography, an application sensitive to geometric errors (Irfanoglu et al. 2012). Thus, the loss of geometric fidelity at 7T undermines the very precision that high-resolution dMRI aims to achieve, reducing the reliability and reproducibility of results across studies.

To improve geometric fidelity in regions impacted by local field inhomogeneities, early diffusion MRI studies relied on field maps to estimate off-resonance effects, or applied nonlinear registration of diffusion images to an undistorted structural scan. These strategies provided partial improvements but were limited by imperfect field estimation or contrast mismatches across modalities. More recently, reversed phase-encoding (blip-up/blip-down) acquisitions have become the standard approach for distortion correction. By acquiring images with opposite phase-encoding polarities – e.g., Anterior-to-Posterior (AP) and Posterior-to-Anterior (PA) – distortions are induced in equal and opposite directions. This allows for algorithms (such as FSLs *topup (Jenkinson et al. 2012; Andersson, Skare, and Ashburner 2003), DR BUDDI* within Tortoise (Irfanoglu et al. 2015, 2024), or *HySCO* within the ACID toolbox (Ruthotto et al. 2013)) to estimate the true displacement field and recover both geometry and signal intensities.

However, the implementation of this approach of using reverse phase-encoding scans for distortion correction varies across studies. Large-scale studies such as UK Biobank (Sudlow et al. 2015) and ABCD (Karcher and Barch 2021) commonly use a single reversed b=0 volume to drive this correction. This single b=0 approach lacks volume-to-volume information and applies a static correction; however, although the main distortion field is static, subject motion causes the *effect* of this field to change from one volume to the next. Others have leveraged full 2-way forward and reverse acquisitions (Left-to-Right (LR) and Right-to-Left (RL)) across all diffusion weighted images (DWIs), as in the Human Connectome Project (Van Essen et al. 2013). This approach can result in lengthy scan times, limiting its potential for widespread adoption. The Developing HCP (Edwards et al. 2022) has addressed these issues in 3T imaging by employing a more time-efficient acquisition, interleaving all four phase-encoding directions (AP, PA, RL, and LR) across non-repeated DWI directions – i.e., a full 4-way acquisition (Hutter et al. 2018; Bhushan et al. 2014).

Systematic evaluations at 3T have confirmed the benefits of these richer acquisition strategies: Irfanoglu et al. (Irfanoglu et al. 2021) demonstrated that correction using full sets of reverse PE DWIs substantially reduces scan-rescan variability of diffusion metrics over single b=0 approaches, and that a 4-way scheme provides the highest reproducibility while further mitigating ghosting and other EPI artifacts. Thai et al. (Thai et al. 2025) extended these findings, showing that multi-directional phase-encoding also yields more anatomically faithful diffusion maps by minimizing residual distortions, PE-direction dependent ghosting and chemical shift artifacts, as well as co-registration errors.

These advantages of multi-phase encoding or reverse phase encoding acquisitions at 3T raise the question of how these strategies perform at 7T, where they may be even more crucial as the magnitude and complexity of susceptibility distortions are greater at higher fields, and the push for sub-millimeter resolution makes geometric fidelity and reproducibility a necessity. Moreover, the powerful gradients that enable this sub-millimeter resolution can also induce stronger eddy current distortions, placing an even greater demand on achieving precise geometric accuracy. In this study, therefore, we sought to systematically evaluate a range of phase-encoding designs to determine their impact on geometric accuracy, SNR, and reproducibility at 7T.

Inspired by Irfanoglu et al. (Irfanoglu et al. 2021), we implemented nine corrected acquisition strategies, each designed to fit within a practical 10-minute scan time at 0.9 mm isotropic resolution but differing in the subsets of phase-encoding directions acquired and the subsequent distortion-correction approaches applied. We then compare these nine corrected acquisition strategies, along with two uncorrected strategies (i.e., without reverse phase-encoding) on multiple metrics of data quality.

As a final validation, we assessed the ability of the best-performing pipeline to reconstruct fine-scale, short-range anatomical pathways using fiber tractography. Short, superficial white matter pathways (often referred to as U-fibers) are small, intricate connections that reside immediately adjacent to the gray-white matter boundary. Because even small distortions can cause misalignment and failure of tractography in these regions, they act as an exemplar demonstration of the power of geometrically accurate high-resolution dMRI.

In sum, this investigation aims to identify acquisition schemes that provide the most faithful alignment with undistorted structural images and the highest reproducibility of derived microstructural measures, while remaining feasible for widespread use in research and clinical settings.

## 2. Methods

Our methodological aim was to design an experiment to test and evaluate nine unique corrected acquisition strategies, each equivalent to ∼10 minute scan time, acquired at 0.9mm isotropic resolution, but with different subsets of phase encoding directions and subsequent distortion correction strategies. To do this, we acquired highly oversampled datasets (Data Acquisition), from which nine time-equivalent corrected acquisitions can be created (Time-Equivalent Acquisitions), along with two uncorrected acquisitions. Broadly, we compared five high-level groups of acquisition and correction approaches: (1) no reverse phase-encoding, (2) single reverse b=0 designs, (3-4) full dual-direction (2-way) datasets and their variants, and (5) 4-way phase-encoding acquisitions. Images were acquired on N=5 subjects, each scanned twice for test-retest evaluation. Each strategy was assessed for distortion correction efficacy, SNR, and scan-rescan reproducibility.

### 2.1 Data Acquisition

DWI data were acquired on the NexGen 7T (Siemens Healthcare), which includes the PNS-optimized Impulse head gradient coil for a Gmax of 200 mT/m and max slew rate of 900 T/m/s [4]. Subjects were scanned using an 8-channel pTx, 64-channel Rx array. Acquisition parameters for 0.9 mm DWI protocol: 152 axial slices, GRAPPA3, MB2, PF6/8, TE=48.4ms, TR∼5500ms, b=1000 s/mm^2^.

The full, highly oversampled acquisition included a 64 diffusion-weighted direction dataset repeated four times, with phase encoding in the AP, PA, LR, and RL directions. These 64 directions were uniformly distributed over a sphere, and sampled so that any subset of N<64 directions is as uniform as possible (or quasi-uniform). A b=0 image was interspersed every 16 volumes. In total, this protocol consisted of four b=0 volumes and 64 diffusion weighted volumes, and took approximately 10 minutes per PE direction, resulting in 40 minutes of DWI acquisition.

### 2.2 Time-equivalent acquisitions

From this oversampled acquisition, we created nine time-equivalent acquisitions (**Figure 1, bottom**), in addition to two uncorrected/unprocessed acquisitions that serve as baseline measurements.

**Figure 1.**
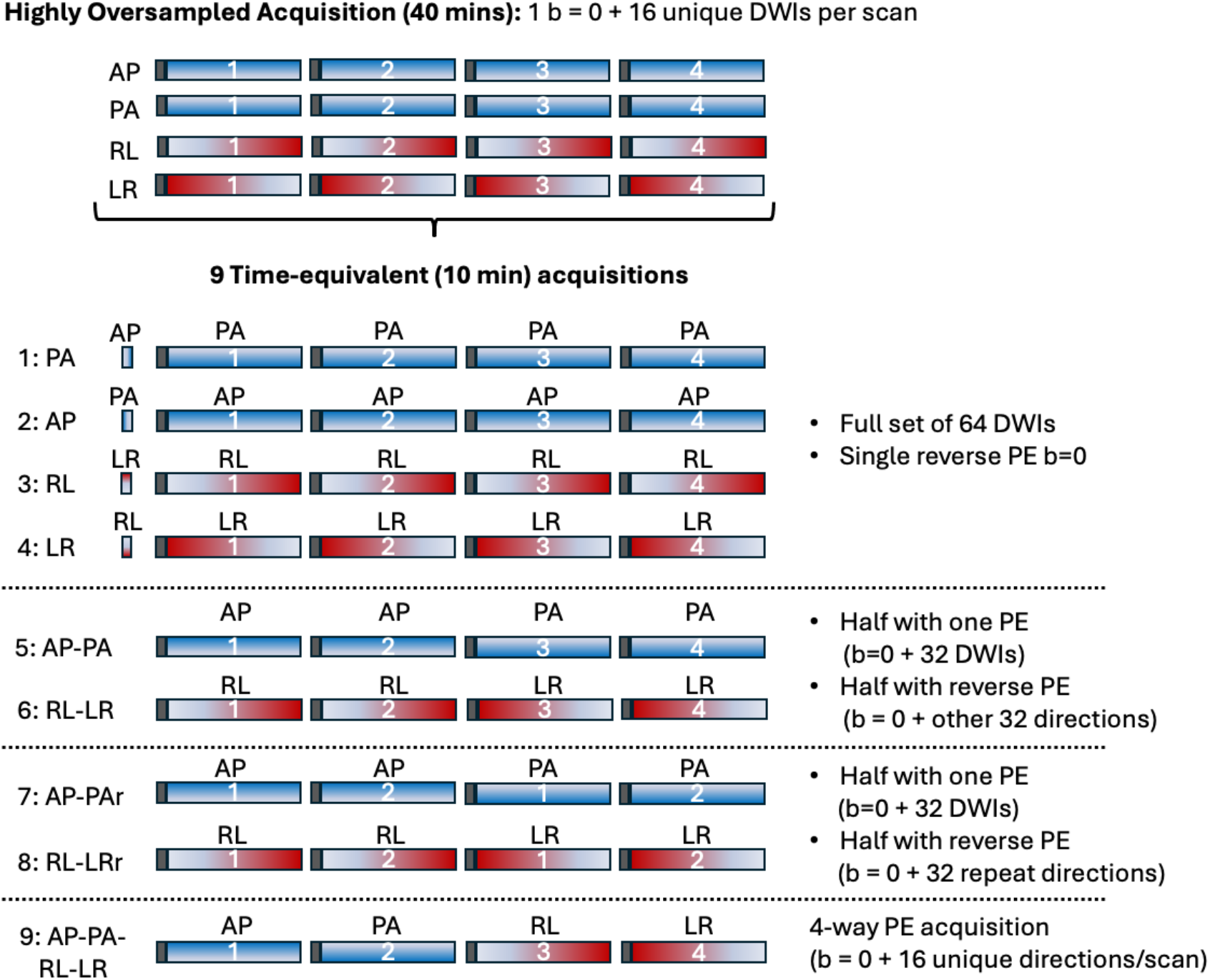
Methodology. The highly oversampled acquisition (top) enabled creation of subset combinations of nine time-equivalent 10 minute acquisitions (bottom). A full 10 min acquisition includes 64 uniformly distributed diffusion weighted directions (with b=0 images interspersed every 16 volumes). This is repeated once for each of the four PE directions (AP, PA, RL, and LR). The nine corrected acquisitions depicted here fall into four categories: (1-4) single reverse PE b=0 (PA, AP, RL, LR); (5-6) full blip-up/blip-down with unique DWIs (AP-PA, RL-LR); (7-8) full blip-up/blip-down with repeated DWIs (AP-PAr, RL-LRr); (9) -way PE (AP-PA-RL-LR).

The two baselines represented the raw acquired datasets that are uncorrected for distortion:

- PAu: uncorrected acquisition with PA PE
- LRu: uncorrected acquisition with LR PE

Acquisitions 1-4 represent common acquisition schemes with a full set of 64 (unique) DWIs with a given PE direction and a single reverse PE b=0:

1. PA: full set of 64 PA DWIs, single reverse AP b=0
2. AP: full set of 64 AP DWIs, single reverse PA b=0
3. LR: full set of 64 LR DWIs, single reverse RL b=0
4. RL: full set of 64 RL DWIs, single reverse LR b=0 Acquisitions 5-6 represent acquiring half the scan (b=0 and DWIs) with one PE, and the other half of the scan with the reverse PE (totaling 64 unique DWIs):
5. AP-PA: half with AP b=0 + 32 DWIs, half with PA b=0 + 32 DWIs
6. RL-LR: half with RL b=0 + 32 DWIs, half with LR b=0 + 32 DWIs Acquisitions 7-8 represent a unique case where we take advantage of FSL eddy algorithm least squares reconstruction. As stated in FSL documentation, this method attempts to use complementary information in images acquired with opposing PE-directions, where a compressed area in one of the images will be stretched in the other image. This method, which enables recovery of spatial resolution in geometrically compressed regions, can only be used if all acquisitions have been repeated with opposed PE-directions [5]. Thus, we acquired half the scan (b=0 and DWIs) with one PE, and the other half of the scan with reverse PE but the exact same DWI sampling (totaling 32 unique DWIs):
7. AP-PAr (“repeated”): half with AP b=0 + 32 DWIs, half with PA b=0 + 32 DWIs
8. RL-LRr (“repeated”): half with RL b=0 + 32 DWIs, half with LR b=0 + 32 DWIs Finally, acquisition nine represents a full 4-way acquisition, in which each quarter of the scan has a different PE direction.
9. AP-PA-LR-RL: 4-way phase encoding with b=0 + 16 DWIs per PE direction

### 2.3 Image preprocessing

Mrtrix3 functions (*mrconvert, mrcat*) were used to extract subsets of each acquisition and concatenate data, creating the nine time-equivalent corrected acquisitions and two (also time-matched) uncorrected acquisitions, for a total of 11 acquisition strategies. It is important to note that all time-equivalent acquisitions were processed separately, after subsets of the oversampled acquisition were created. Each acquisition was denoised (*Patch2Self* algorithm) and corrected for susceptibility distortions, motion, and eddy currents using the FSL software *topup* and *eddy* algorithms. Diffusion tensors were fit using linear least squares (dtifit, FSL). T1-weighted images were skull stripped (*hd-bet*) and white matter, gray matter, and CSF was segmented using FSL’s *fast*. Finally, diffusion data was aligned to T1 contrast using epi_reg (FSL), enabling comparison of contrasts in either diffusion or T1 spaces.

### 2.4 Image analysis

Each of the 11 time-equivalent acquisition strategies was evaluated to identify optimal acquisition and preprocessing schemes. An optimal strategy was defined by its ability to deliver: (1) high geometrical alignment with an undistorted structural image, (2) high SNR, and (3) high scan-rescan reproducibility of derived diffusion measures.

Diffusion tensors were estimated at each voxel with FSL’s dtifit, which fits a tensor via linear least-squares. From the fitted tensors we derived fractional anisotropy (FA), mean diffusivity (MD), axial diffusivity (AD = λ_1_), and radial diffusivity (RD = (λ_2_+λ_3_)/2). SNR was computed as the ratio of the mean diffusion signal to the residual standard deviation from the tensor fit (σ = √[SSE/DOF]) and then averaged within a white-matter mask - using the fit residuals as a proxy for a per-voxel noise estimate.

Geometric accuracy was assessed both qualitatively and quantitatively. Qualitatively, anatomical boundaries derived from the high-resolution T1-weighted image (e.g., the brain perimeter and the gray-white matter interface) were overlaid onto the processed diffusion images (*b*=0) and DTI-derived maps (FA, MD, AD, RD) for visual inspection of alignment. Quantitatively, Mutual Information (MI) between the subject’s T1-weighted image and the corresponding *b*=0 and FA maps were calculated. A higher MI value indicates better geometric alignment between the modalities. Mean absolute difference in image gradient magnitude between the normalized T1 and *b*=0 images were also calculated (i.e., Gradient Difference; GD), where a higher value indicates greater alignment of image boundaries/structures.

Scan-rescan reproducibility was assessed using the two separate imaging sessions (acquired on different days) for each participant. We calculated two key metrics for the DTI-derived maps (FA, MD, AD, RD): (1) MI between the maps from scan 1 and scan 2, where higher MI signifies greater reproducibility; (2) Mean Absolute Error (MAE) between the maps from scan 1 and scan 2, where a lower MAE indicates better reproducibility.

Analyses were performed across the whole brain to assess general performance and also within the anterior lateral prefrontal cortex (aLPFC) to demonstrate performance within a targeted region of interest for studies exploring short-range white matter connectivity. Additionally, we assessed the ability of the best-performing pipeline to reconstruct fine-scale anatomical pathways using fiber tractography. This application was chosen as it serves as a comprehensive test of data quality, requiring high SNR, high-resolution, and precise geometric alignment with the gray-white matter boundary for accurate seeding and tracking. The ability to resolve short, superficial white matter association fibers served as a key benchmark for pipeline performance.

### 2.5 Statistical Analysis

Statistical analyses were performed in MATLAB to compare the performance of the 11 dMRI acquisition and correction strategies. Given the repeated-measures design, the non-parametric Friedman test was chosen as the primary statistical tool. A significance level of p < 0.05 was used for all tests. The analysis was structured as a three-step process for each metric (e.g., MI, GD, MAE).

For analysis, the 11 strategies were first grouped into five high-level strategies based on their acquisition and correction approach: (1) uncorrected baseline with single phase encoding direction (PAu, LRu); (2) single reverse PE b=0 (PA, AP, RL, LR); (3) full blip-up/blip-down with unique DWIs (AP-PA, RL-LR); (4) full blip-up/blip-down with repeated DWIs (AP-PAr, RL-LRr); (5) 4-way PE (AP-PA-RL-LR).

Tests were performed for:

1. Between-Group Differences: To test for differences between the five high-level strategies, we first averaged the metric scores for conditions within the same group for each subject. A Friedman test was then performed on these five group-level averages. If the result was significant, post-hoc multiple comparison tests were used to identify which groups were statistically different from one another.
2. Within-Group Differences: To determine if the choice of specific PE direction (e.g., AP vs PA) mattered within a group, a separate Friedman test was conducted for each multi-member group (Groups 1-4). A significant result indicated meaningful performance differences between strategies within that category.
3. Identification of Top-Performing Strategies: To identify the “best” individual strategies, we first found the condition with the optimal median value for a given metric (highest for accuracy, lowest for error). Then, every other strategy was compared directly to this “best” one using a Wilcoxon signed-rank test. Any strategy not significantly different from the best (p > 0.05) was considered statistically equivalent and included in the “top-performing tier.”

Together, this enabled evaluating both high-level acquisition strategies and specific phase-encoding implementations and identification of optimal acquisition and correction strategies.

## 3. Results

### 3.1 Geometrical Accuracy and SNR

Qualitative assessment shows that distortion correction improves geometrical accuracy (**Figure 2**). In the uncorrected strategies (PAu, LRu), misalignment between the FA maps and anatomical boundaries is evident upon visual inspection, especially in the prefrontal and temporal regions. In contrast, all nine correction strategies show improved alignment, with the FA and color-coded FA maps matching the underlying anatomical structures from the T1-weighted image.

**Figure 2.**
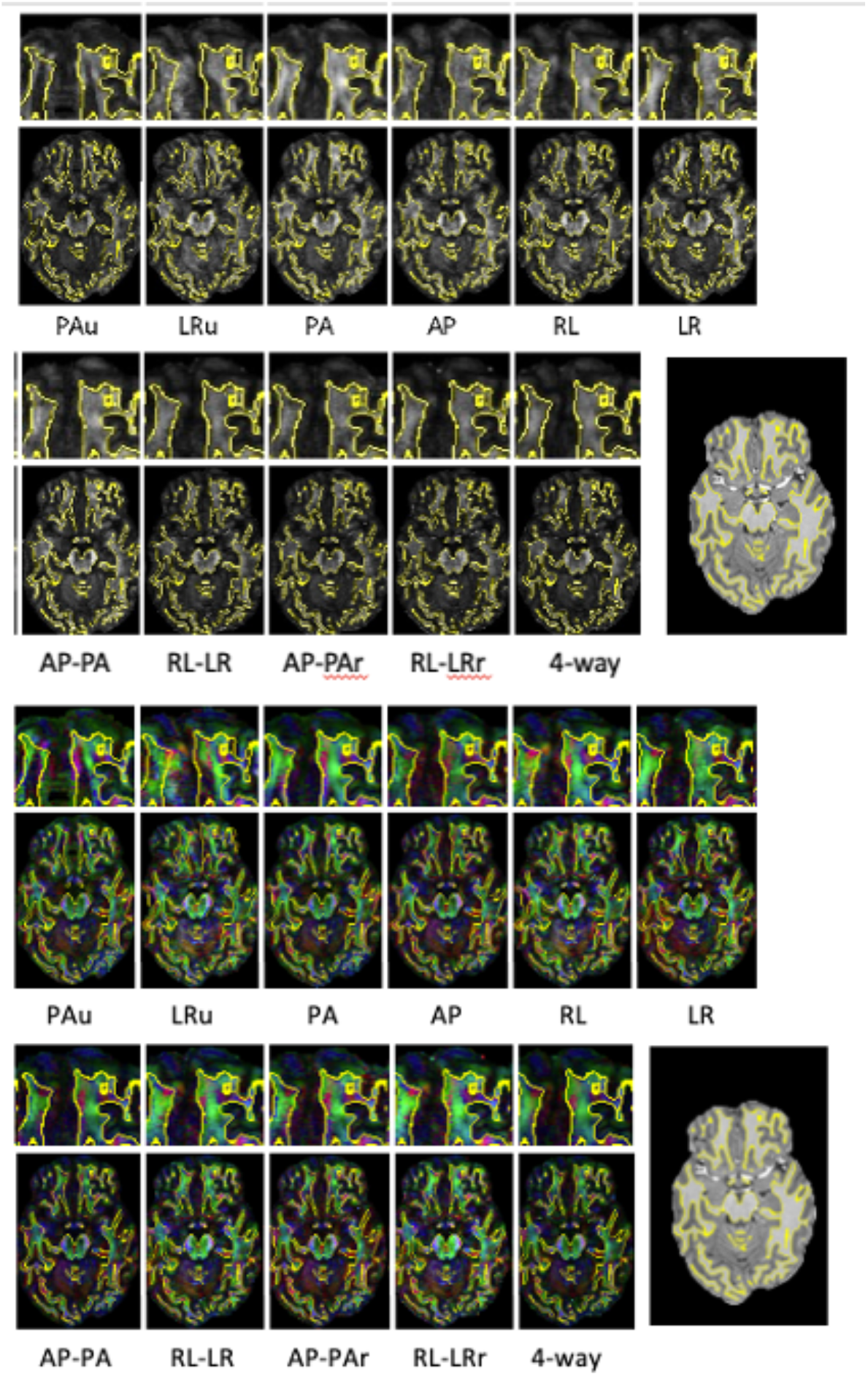
Qualitative alignment of FA with T1. Fractional Anisotropy (FA) maps (top) and color-coded FA maps (bottom) are shown for all 11 strategies for a single representative subject. Anatomical boundaries derived from the T1-weighted image are overlaid in yellow. The two uncorrected strategies (PAu, LRu) show clear geometric distortions, while all correction strategies demonstrate improved alignment with small visual differences in anisotropy and orientation.

This is confirmed quantitatively (**Figure 3**), where the uncorrected strategies performed worst across all geometrical accuracy metrics. The top-performing tier included those strategies with a full blip-up/blip-down acquisition (AP-PA, RL-LR), a repeated acquisition (AP-PAr, RL-LRr), and the 4way PE acquisition (AP-PA-RL-LR) – i.e., strategies 5-9 in Figure 1. These methods had significantly higher MI and lower GD than uncorrected data and generally outperformed those that use only a single reverse PE b=0 image. While these strategies were often statistically equivalent, the 4-way strategy frequently had the highest median performance. In SNR analysis, the “repeated” acquisitions had a significantly higher SNR than all other strategies; this is an expected result, as the preprocessing for these acquisitions averaged the two identical sets of diffusion-weighted images during motion/eddy-current correction, thereby reducing noise. Differences in SNR among the other strategies were not statistically significant.

**Figure 3.**
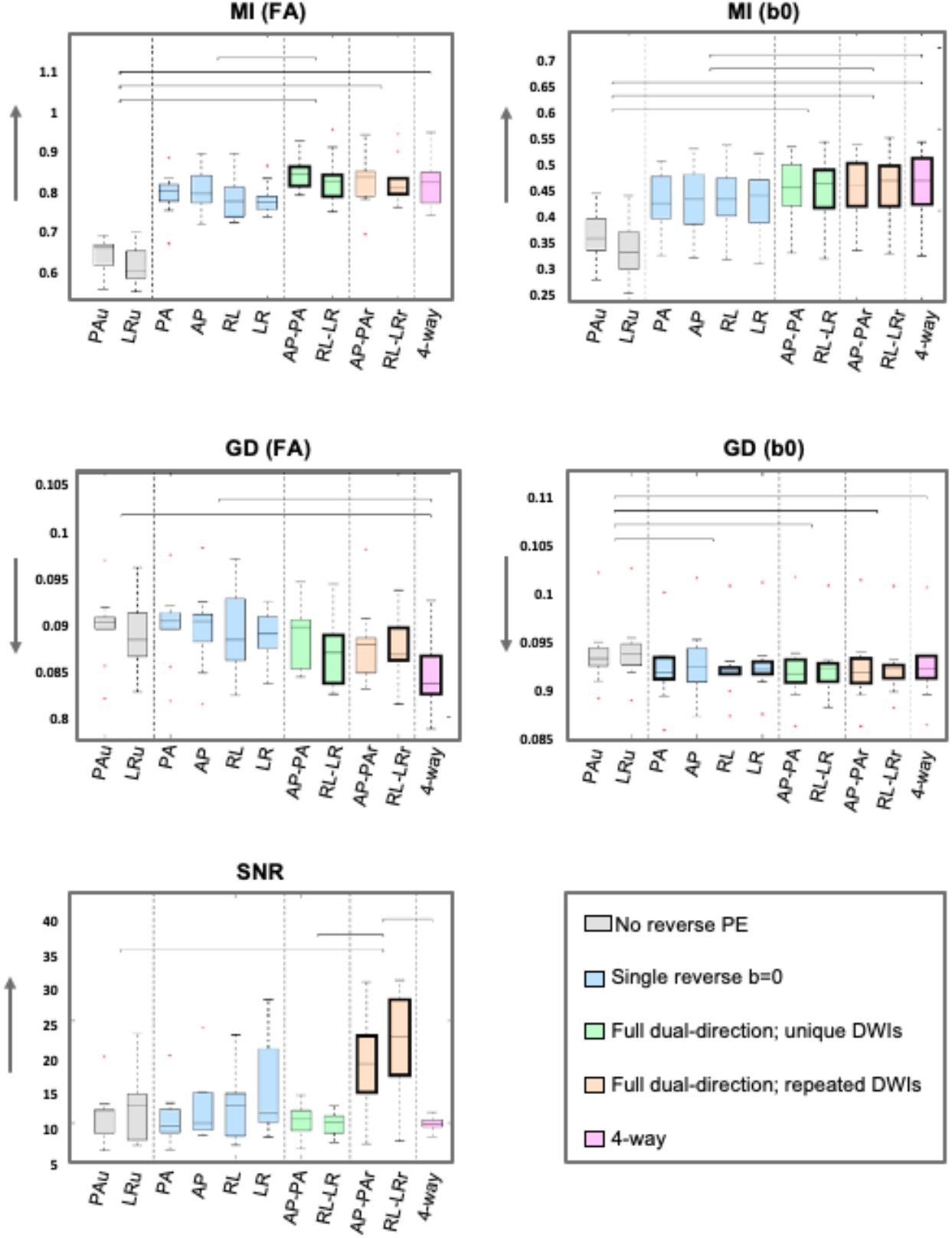
Quantitative metrics for Geometrical Accuracy and SNR. Boxplots show the distribution of MI, GD, and SNR across all subjects for the 11 strategies. Arrows indicate the direction of improvement. Bolded boxes highlight the top-performing tier of strategies that are not statistically different from the single best performer for each metric. Lines above the plots indicate statistically significant differences between groups. Significant within-group differences were observed between PAu and LRu in MI (b0), and between AP and RL in GD (b0).

### 3.2 Reproducibility

Qualitative maps of scan-rescan error, created by averaging the absolute difference between sessions across all subjects, are shown in **Figure 4** for both FA and RD. While whole-brain improvements in reproducibility across the tested correction strategies are subtle, a general trend shows voxel-wise errors decrease when moving from uncorrected (left) to the more comprehensive correction schemes (right). This improvement is most visible in the 4-way acquisition, which shows a widespread reduction in differences – particularly for RD.

**Figure 4.**
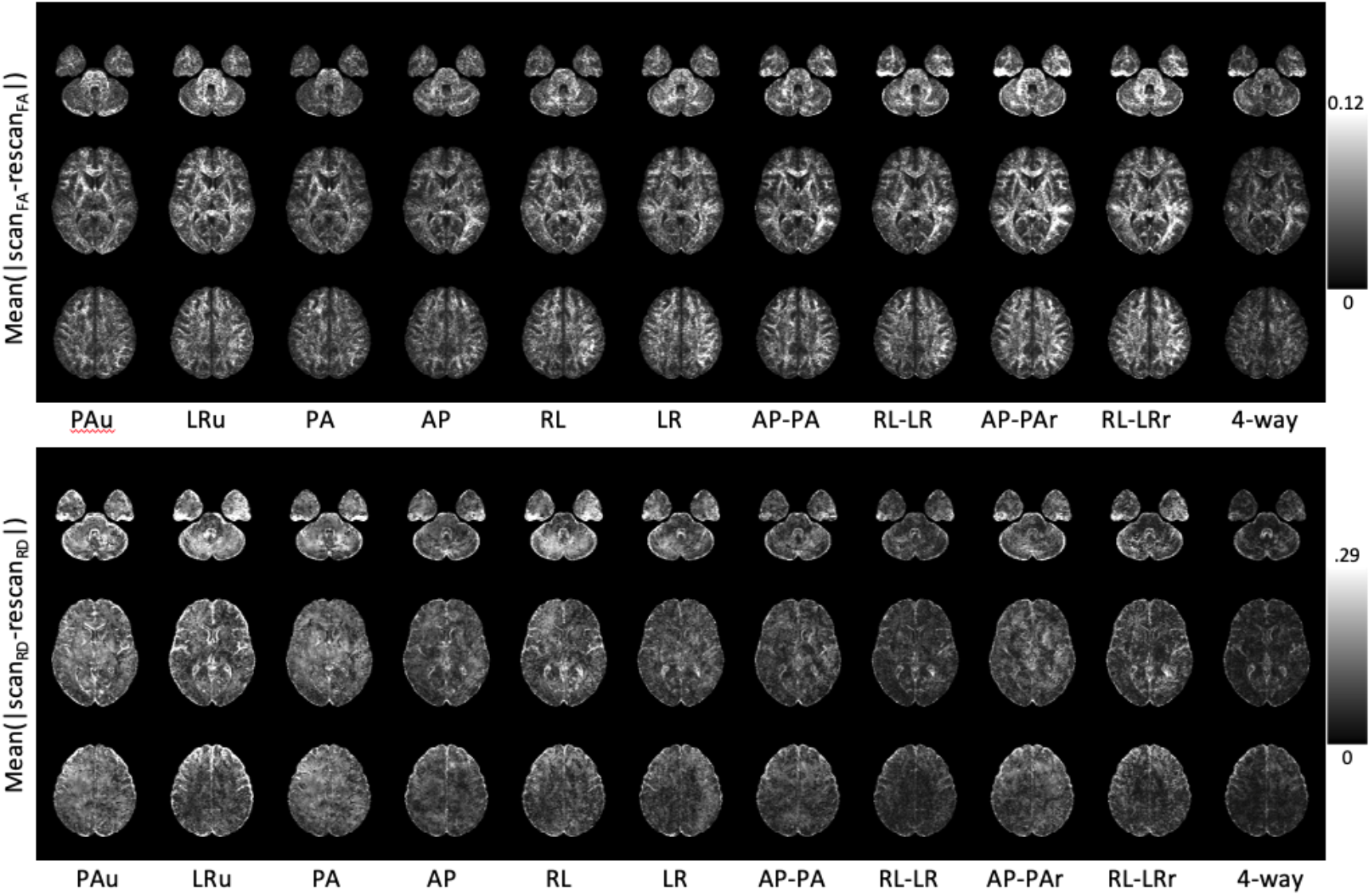
Qualitative maps of scan-rescan reproducibility. The mean absolute difference between scan 1 and scan 2, averaged across all subjects, is displayed for FA (top) and RD (bottom) for all 11 strategies. Brighter voxels indicate higher scan-rescan error (i.e., lower reproducibility).

The quantitative analysis of reproducibility (**Figure 5**) generally aligns with the findings for geometrical accuracy. Across all metrics, the uncorrected strategies (PAu, LRu) performed significantly worse than all corrected strategies, showing the lowest MI and highest MAE. Among the corrected methods, the 4-way PE strategy generally outperformed the full blip-up/blip-down acquisitions, which in turn outperformed the single reverse b=0 methods. However, due to inter-subject variability, these differences were not statistically significant. Despite this, the 4way PE acquisition (AP-PA-RL-LR) consistently demonstrated the most optimal median performance, achieving the highest MI and the lowest MAE for both FA and RD.

**Figure 5.**
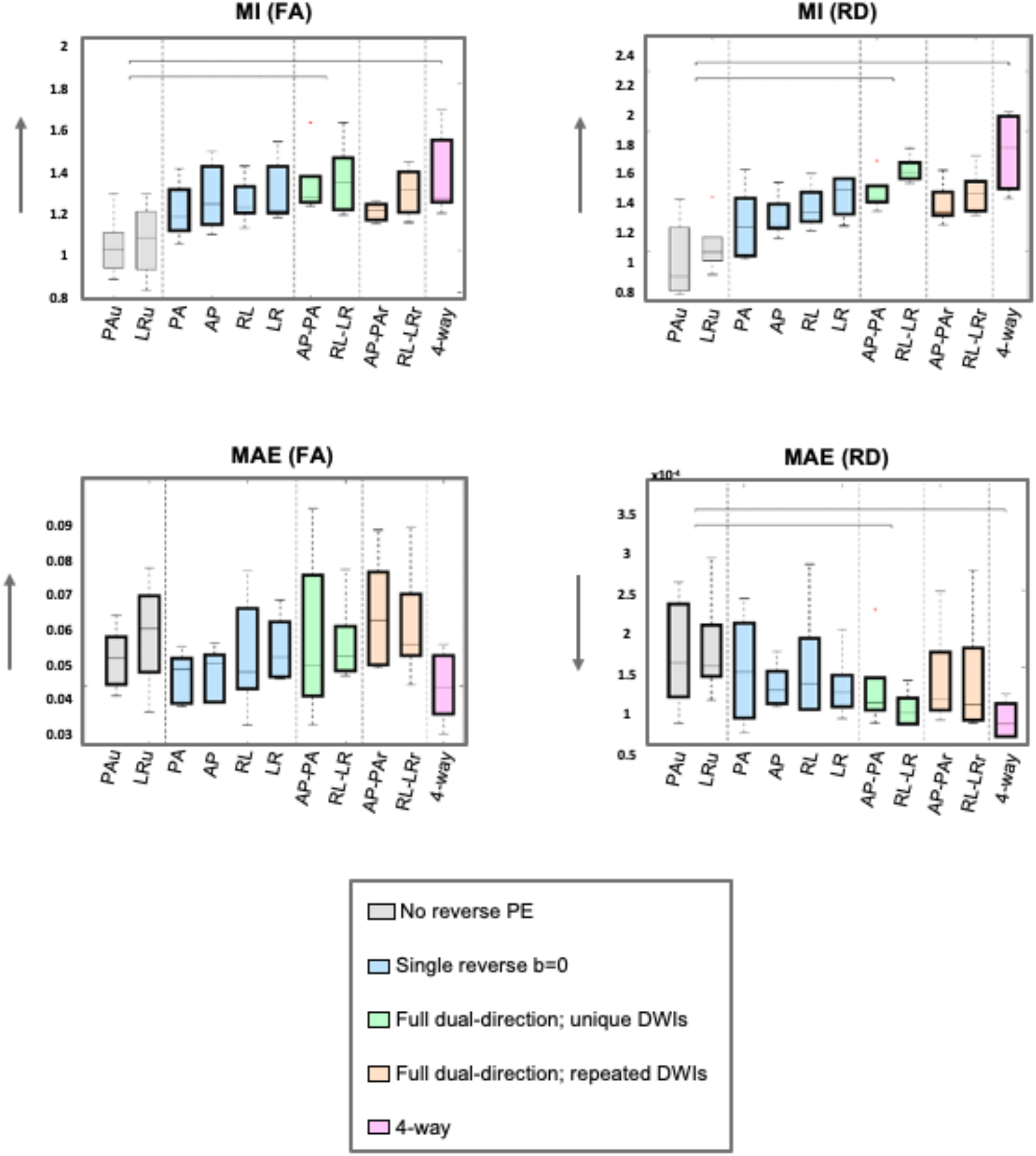
Quantitative metrics for scan-rescan reproducibility. Boxplots show the distribution of MI and Mean MAE between scan 1 and scan 2 for FA and RD across all subjects. Arrows indicate the direction of improvement. Bolded boxes highlight the top-performing tier of strategies. Lines above the plots indicate significant differences between groups, and asterisks below indicate significant differences within a group (none indicated in these tests).

### 3.3 Region-Specific Analysis and Validation with Tractography

To test if the whole-brain findings generalize, we repeated the quantitative analysis of geometrical accuracy and SNR within the targeted aLPFC region of interest (**Figure 6**). The results were generally consistent with the whole-brain analysis. The uncorrected strategies again performed the worst, while the top-performing tier consistently included the full blip-up/blip-down (both unique and repeated), although not always the 4-way PE acquisition. We observed greater variability across the metrics, as expected for a smaller ROI. This regional consistency suggests the benefits of comprehensive correction strategies are not driven solely by large-scale effects (contrast changes in CSF/brainstem) but extend to localized cortical structures.

**Figure 6.**
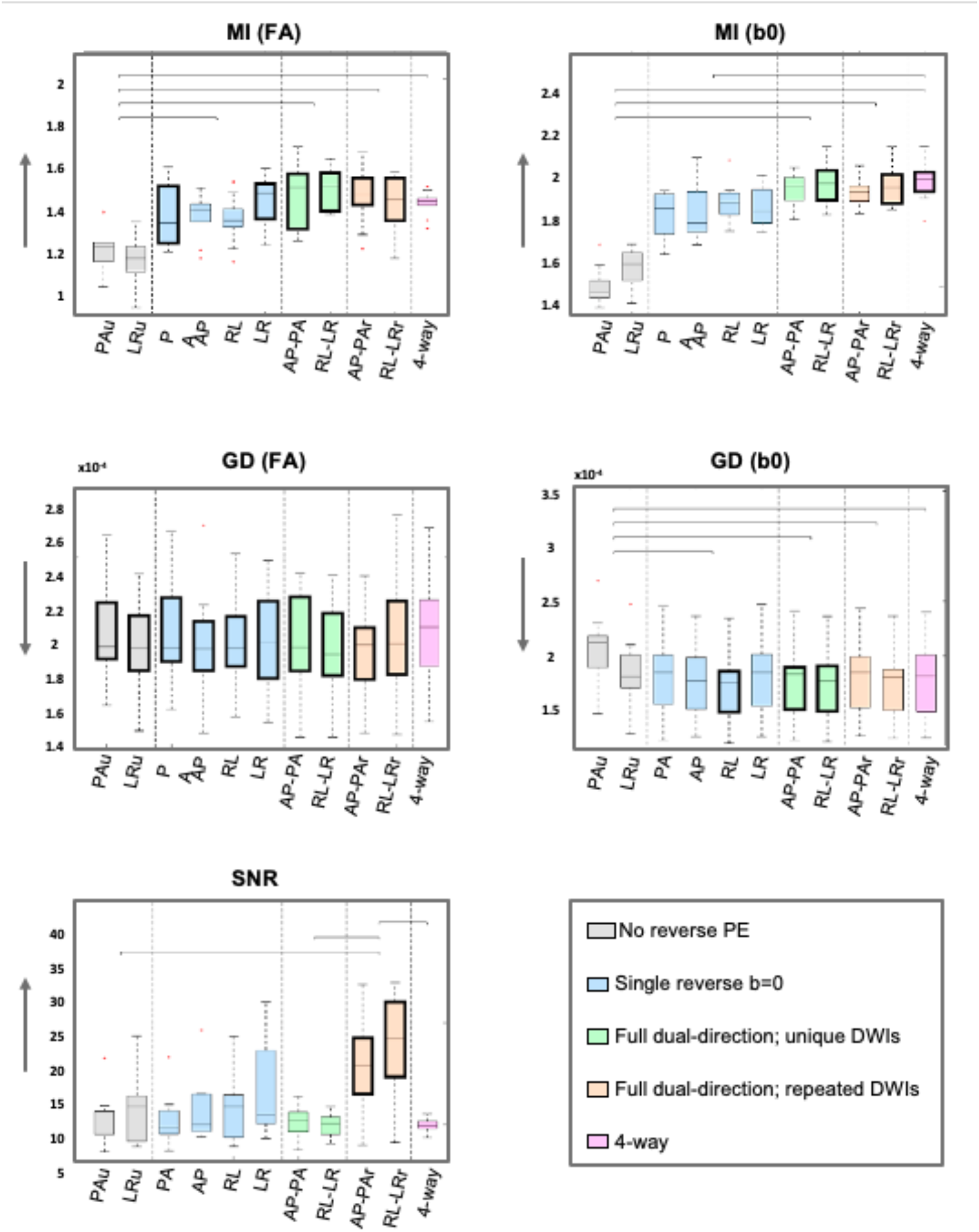
Quantitative metrics within the Prefrontal Cortex ROI. Boxplots show the distribution of MI, GD, and SNR calculated within the aLPFC for all 11 strategies. Results largely mirror the whole-brain analysis, with arrows indicating the direction of improvement and bold boxes highlighting the top-performing tier. Lines above the plots indicate significant differences between groups. Significant within-group differences were observed between Pau and LRu in GD (b0), between AP and RL in GD (b0), and between RL and LRr in GD (b0).

Finally, to demonstrate the practical benefits of an optimized acquisition, **Figure 7** showcases the high-quality data and tractography results from the top-performing 4-way PE strategy. The top row illustrates the high image quality: raw DWIs have high SNR with clear gray-white matter boundary, and the derived FA and color-coded FA maps show fine anatomical detail and coherent orientation patterns. The bottom row demonstrates the utility of this data for advanced tractography applications. As expected, major long-range white matter pathways are robustly reconstructed (bottom left). Importantly, this acquisition enables the ability to resolve fine-scale, short-range superficial white matter pathways, which form a thin sheet immediately adjacent to the cortex (bottom right). This level of detail enables the in vivo virtual dissection of connections between specific cortical regions that have strong histological evidence but have been challenging to validate in living humans due to limitations of conventional imaging (bottom right; inset).

**Figure 7.**
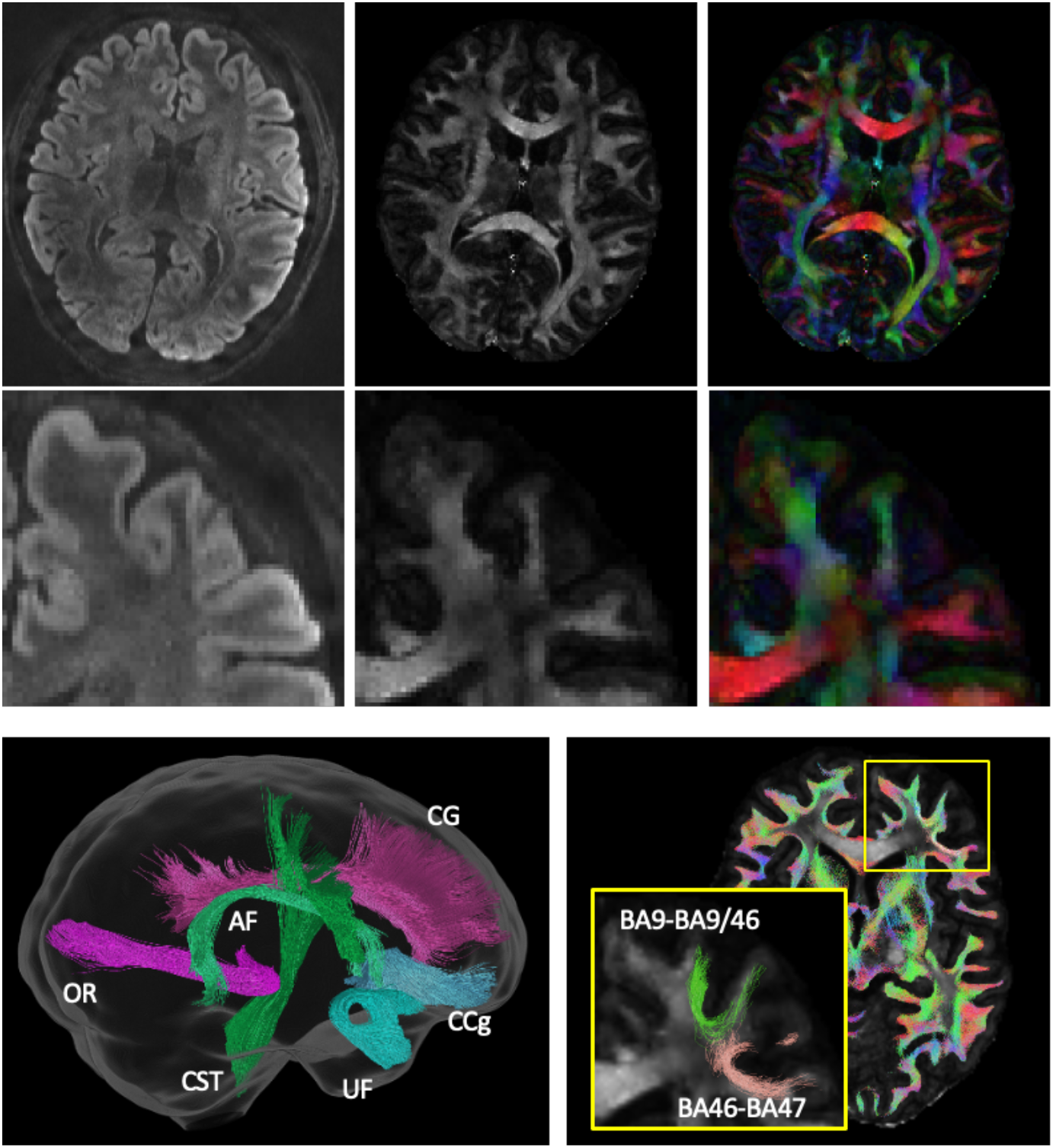
Qualitative results and tractography from the optimal 4-way PE acquisition. Top row: a representative diffusion-weighted image, FA map, and color-coded FA map demonstrate high SNR and fine anatomical detail. Bottom row: reconstructed long range pathways (left; Arcuate Fasciculus, Cingulum Bundle, genu of Corpus Callosum, Uncinate Fasciculus, Corticospinal Tract, Optic Radiations) and short-range superficial white matter pathways (right). Inset highlights the ability to resolve exemplar fine-scale connections between histologically defined Brodmann areas (BA9-BA9/46 and BA46-BA47).

## 4. Discussion

In this study, we systematically evaluated a range of time-matched acquisition and correction strategies for high-resolution 7T diffusion MRI. Our results on image quality and reproducibility show a clear hierarchy: 1) implementing any form of distortion correction improves geometric accuracy over uncorrected data; 2) a full set of reverse PE DWIs – either 2-way or 4-way – results in higher geometric accuracy and reproducibility than using only a single reverse b=0 image; and 3) 4-way PE consistently provided the highest image fidelity. These findings at 7T confirm and extend the principles established at 3T (Irfanoglu et al. 2012, 2021), validating them in a more challenging ultra-high-field context, in which artifact correction is critical.

### 4.1 Value in full DWI acquisitions

Our results confirm the intuitive finding that distortion correction is necessary to improve geometric accuracy. More importantly, this work enables insight into specific benefits of more comprehensive acquisition and correction strategies. We found that acquiring a full set of reverse PE DWIs—either 2-way or 4-way—is generally better than relying on a single reverse b=0 pair, which is likely the most common approach used in research-based applications. This advantage arises because the full set of reverse DWIs provides more complete information to the eddy correction tool. As noted previously, while the main distortion field is static, subject motion causes the effect of the distortion field to change from one volume to the next, thus the single b=0 correction approach lacks volume-to-volume information (i.e. for each individual DWI volume), applying a static correction, which may be less robust over longer scan periods given that subject motion causes the effect of the main distortion field to change across volumes. By including both blip-up and blip-down data for all DWIs, eddy (or any tool that models distortion, motion, and eddy currents) can more accurately model these dynamic changes and separate them from true (or estimated) head motion.

A separate consideration is the comparison of unique DWIs directions versus repeated DWI directions – i.e., AP-PA vs. AP-PAr and RL-LR vs. RL-LRr. The repeated acquisition enables a specific “least-squares reconstruction” feature in eddy (Andersson, Skare, and Ashburner 2003) (in contrast to the default Jacobian-based resampling) that simultaneously models the “true” undistorted image from the two oppositely distorted (compressed/stretched) acquisitions, and can theoretically recover signal and lost resolution in areas that have been compressed or stretched (Andersson, Skare, and Ashburner 2003). However, we found little *consistent* differences in geometric accuracy between the unique and repeated strategies, where the most significant difference was an increase in SNR for the repeated acquisition – an expected outcome of averaging two identical measurements. That said, the “optimal” acquisition will almost certainly depend on the subsequent use: microstructure modeling, orientation estimations, tractography reconstruction, etc.

Further, our results show that 4-way acquisition offers advantages in image fidelity and reproducibility over even full 2-way schemes. This improvement is likely attributable to mitigation of EPI artifacts beyond primary distortion correction, for example N/2 Nyquist ghosts. As shown previously (Thai et al. 2025; Huber et al. 2025), ghosting and other eddy current-related artifacts are PE-direction (and readout-direction) dependent, whereby a ghost that contaminates a region in an AP acquisition, for example, may be displaced outside the brain in an LR acquisition. By acquiring and averaging data across four directions, these spatially inconsistent artifacts are suppressed. This artifact mitigation is especially valuable at 7T, where all artifacts are exacerbated.

Finally, our short-range tractography results validate that the 4-way acquisition offers the anatomical accuracy necessary to make these challenging reconstructions possible in vivo. By minimizing error, we can precisely define seeding, ending, and inclusion/exclusion masks that enable anatomically accurate reconstruction of these pathways.

These findings should be interpreted with several caveats in mind. First, they are based on a limited sample size, with only two repeats. Second, what can be considered a “DTI” acquisition, although it is high angular resolution, does not include multiple shells or different encodings. Third, NexGen 7T - hardware, but generally applicable to any high-resolution imaging at any field or with any hardware and aligns with results of low res 3T.

### 4.2 Recommendations and Future Directions

While further research is needed to determine whether the present results generalize to multiple shells, different encodings, or lower-resolution scanners, we offer practical recommendations as a starting-point for researchers designing high-resolution 7T dMRI studies.

1. For highest fidelity, 4-way encoding provides the best overall performance, yielding the highest geometric accuracy, artifact suppression, and reproducibility. We acknowledge, however, that implementing this scheme can be challenging, as it may require careful optimization to match parameters (field-of-view, resolution, bandwidth, TE/TR) across both AP/PA and LR/RL acquisitions. Furthermore, differences in PNS limits on different readout axes (Feinberg et al. 2025) may prevent parameter matching depending on the desired protocol.
2. Acquiring a full set of 2-way DWIs is an excellent alternative that provides most of the benefits of the 4-way acquisition. While we do not explicitly study the effects of number of gradient directions here, software documentation suggests that benefits of repeated acquisitions (least-squares reconstruction) might be more beneficial once 60+ directions are repeated (FSL eddy; FSL v5.0.10 documentation: https://web.mit.edu/fsl_v5.0.10/fsl/doc/wiki/eddy.html).
3. While it quantitatively underperformed more comprehensive schemes, the common approach of using a single reverse b=0 pair provides significant improvement over uncorrected data on the metrics of geometric accuracy, artifact suppression, and reproducibility. Given its minimal time cost, it remains a pragmatic choice when acquiring diffusion data.

Finally, beyond PE strategy, we echo several general principles for high quality diffusion MRI acquisition from previous studies (Irfanoglu et al. 2012, 2021; Thai et al. 2025): maintain high angular resolution with sufficient unique directions to properly model complex fiber orientations (Jones, Horsfield, and Simmons 1999; Tournier, Calamante, and Connelly 2013); sample gradients over the entire sphere, rather than a hemisphere, to balance the effects of imaging gradients; match imaging parameters (TE, TR, resolution) across all scans to ensure comparable contrast and sensitivity; and use appropriate preprocessing pipelines (Cieslak et al. 2021; Cai et al. 2021; Chen et al. 2024), with software tools (Irfanoglu et al. 2015; Andersson and Sotiropoulos 2016) specifically designed to leverage multi-phase encoding datasets.

## Acknowledgments

Research was supported by grants from the National Institutes of Health: BRAIN Initiative grant U24-NS129949 (D.A.F., A.T.V, A.J.S.B.), R01-MH133637 (S.A.B.), U01-EB025162 (D.A.F., A.T.V, A.J.S.B., R01-MH111444 (D.A.F., A.T.V, A.J.S.B.), R44-MH129278 (D.A.F., A.T.V, A.J.S.B), T32-EB001628 (M.A), and K01-EB032989 (K.G.S.). Additional support was provided by the Chancellor’s Office of UC Berkeley (D.A.F.) and Weill NeuroHub (D.A.F., A.T.V, A.J.S.B).

